# Despotic long-tailed macaques benefit others in a group service paradigm

**DOI:** 10.1101/2024.06.28.601131

**Authors:** E.J.A.M. de Laat, S. Waasdorp, T.S. Roth, J.J.M. Massen, E.H.M. Sterck

## Abstract

For animals living in social groups, cooperation is a key factor to success. It has been postulated that in social systems with cooperative breeding or a tolerant dominance style, individuals will benefit each other. Cooperation is, therefore, not expected in long-tailed macaques, since they do not breed together and experience a steep unidirectional hierarchy. However, previous studies have shown that they can be prosocial in a dyadic setting. This would comply with the more recently postulated dyadic interdependence hypothesis. To be able to compare their cooperative performances with other species, we set up a group service paradigm similar to that, which has been tested in a number of other species. We presented a swing set apparatus, which an actor could pull in the middle to provide a reward to another individual but without access to the reward for the actor, to three groups of socially housed long-tailed macaques. The macaques showed prosocial behaviour in the test significantly more often than in two control condition. They preferably provided to kin. The prosocial behaviour of the despotic, individual breeding long-tailed macaques counters the cooperative breeding and self-domestication hypotheses, yet supports the dyadic interdependence hypothesis, although future studies on other macaque species with more tolerant dominance styles should elucidate the effect of dominance styles on prosociality.

## INTRODUCTION

Prosocial behaviour, acts from an individual that directly reward the receiver but do not benefit the provider (Burkart et al., 2018; Martin et al., 2021; Massen et al., 2011; Melis, 2018), is considered crucial for initiation and maintenance of cooperation. This is an important benefit of living in a social group (primates: Burkart et al., 2014; Callaghan & Corbit, 2018; Silk & House, 2011; humans: Antón et al., 2014; Hill et al., 2009). Originally, it was presumed that only humans could act prosocially (Hare, 2017; Maclean, 2016), however, a multitude of studies showed some form of prosociality in a wide range of primate species (reviewed in Cronin, 2012). Yet several studies on chimpanzees (*Pan troglodites*) and bonobos (*P. paniscus*) have failed to show prosocial tendencies, even though these two ape species are considered very cooperative (Silk et al., 2005; Silk & House, 2011; Tennie et al., 2016; Van Leeuwen et al., 2021; Verspeek, van Leeuwen, Laméris, Staes, & Stevens, 2022; Vonk et al., 2008). This between-species variation in prosocial behaviour has been linked to species differences in social systems, namely depending on cooperative breeding (Burkart et al., 2014) and self-domestication (Hare, 2017), i.e. egalitarian dominance hierarchies.

To understand the evolution of prosociality, comparative studies are required, however the multitude of different experimental paradigms, set-ups and procedures (reviewed in Cronin, 2012; Dale et al., 2016) make such comparisons difficult. In a first attempt to standardize the experimental paradigm and its procedures across a diverse group of species, Burkart and Van Schaik designed a group-service paradigm (Burkart & Van Schaik, 2013). In such an open setting with active partner choice, individuals can voluntarily provide food to their group-members, while they are not able to reach for that food themselves, nor are they rewarded for doing so in any other way. In a multi-lab effort, they subsequently tested this paradigm in 15 different primate species, and found that prosocial behaviour correlates with extensive allomaternal care (Burkart et al., 2014). These findings supported the cooperative breeding hypothesis (Burkart et al., 2009), which suggests that helping others to rear their young requires high social tolerance and prosociality. Accordingly, empirical evidence for a link between prosociality and cooperation is also found in the cooperatively breeding common marmosets, *Callithrix jacchus*, where in an experimental set-up cooperative success increases twofold when there was at least one prosocial individual in the test dyad (Martin et al., 2021). Further support for this hypothesis comes from a recent avian study, i.e., a comparison of eight different corvid species, which also found the highest prosocial tendencies among the cooperatively breeding birds (Horn et al., 2020). The study on corvids, however, also showed that colonially nesting species have relatively high prosocial tendencies, and these authors suggest that heightened social tolerance (at the nest) may have been an evolutionary pathway to prosociality (Horn et al., 2020).

The evolutionary selection pressure towards the most friendly individuals leads to self-domestication (Hare et al., 2012), and prosociality may, therefore, be restricted to the more egalitarian primate species. This subsequently enhances the probability of prosocial behaviour, such as in Tonkean macaques (*Macaca tonkeana*, Joly et al., 2017; Scopa & Palagi, 2016). Interestingly, however, using a similar group-service paradigm as the previous comparative studies (Burkart et al., 2014; Burkart & Van Schaik, 2013; Horn et al., 2020), Bhattacharjee and colleagues (2023) recently showed prosocial tendencies in Japanese macaques, *Macaca fuscata*. This species is considered very despotic (class I: Thierry et al., 2000; Thierry, 2007; 2020), which makes it a less obvious candidate. They argue, however, that social tolerance at the dyadic level may also spark prosocial motivations, particularly when individuals rely on social bonds for alliances to cope with the complexities of their social organisation (Bhattacharjee et al. 2023). This overarching interdependency hypothesis (Tomasello et al. 2012; Bhattacharjee et al. 2023), therefore, suggests that both interdependencies at the dyadic level (e.g. alliances; kinship) as on the group level (cooperative or colonial breeding) may have caused the evolution of prosociality in different taxa. They therefore call for more analyses on the dyadic level, as well as for more studies on species that are considered rather despotic (Bhattacharjee et al., 2023).

The open setting of the group service paradigm allows, due to testing this in a group setting, for more natural interactions between the participants. This, however, can also cause more dominant individuals to ‘pressure’ others into provisioning to them (Horn et al., 2020). Therefore, prosociality can be classified as proactive or reactive. Prosocial behaviour is characterised as reactive when the provider is triggered by an external motivation, like awareness of a signal of need, an audience present at a close social distance (Jaeggi et al., 2010) or a threat of pressure (Horn et al. 2020). Prosocial behaviour is characterised as proactive when a solicitation by a group member is absent and the behaviour originates out of an intrinsic motivation. A high intrinsic prosocial motivation may keep cooperation going on, without immediate reciprocal rewarding (Jaeggi et al., 2010).

Long-tailed macaques, *Macaca fascicularis*, while not as despotic as the Japanese macaques, are also categorized as a despotic species, class II (Adams et al., 2015; Thierry, 2007). Females are philopatric, whereas males disperse at the age of 3-4 years old. Infants inherit the rank under their mother due to their mother’s support, leading to nepotism (Van Noordwijk & Van Schaik, 1985; Van Schaik & Van Noordwijk, 1987). This means that they will favour kin and are less tolerant towards non-related group members. On top of this, these macaques do not help to raise other’s offspring, therefore any prosociality from these macaques would not conform to the cooperative breeding hypothesis (Burkart et al., 2014). Long-tailed macaques eat alone, they do not share food like capuchin monkeys, which is also seen as a prerequisite for prosocial behaviour (Brosnan, 2010), resulting in a unexpected candidate primate species for prosocial behaviour. Nevertheless, a previous study indicated that these macaques do benefit group members in a dyadic behavioural experimental set-up, that dominant individuals are more prosocial than subordinate ones, and that they are most prosocial towards kin (Massen, 2012). Further tests showed that when the long-tailed macaques could choose whom to benefit in a triadic set-up, that they consider the relative dominance position of their partners more than the relationship quality between themselves and the potential receiver (Massen et al., 2011). In a forced choice-task, however, long-tailed macaques prefer to take a high value food reward by themselves, above providing prosocially (Sterck, et al., 2015). This calls for exploring prosocial behaviour in this despotic species in a group setting.

The aim of this study was to test prosocial behaviour of long-tailed macaques with a group service paradigm (Bhattacharjee et al., 2023; Burkart et al., 2014; Burkart & Van Schaik, 2013; Horn et al., 2020; Martin et al., 2021). In line with the previous studies in this species, we expect dominant individuals to participate more often than subordinates. Moreover, we expect that younger animals will participate more often than adults, because they are more playful and explorative (Tan et al., 2018). Because females will stay in their natal group (Van Schaik et al., 1983), and have to cooperate more intensively with group members, we expect them to be more prosocial than males. Alternatively, males may work together because they form male-male coalitions and disperse together (Mishra et al., 2020; Van Noordwijk et al., 1985) and males may provide to females, for example as a strategy to gain mating success (cf. friendships: Massen et al., 2012). In addition, we expect that the individuals that act prosocially will provide kin more than non-kin (Massen et al., 2011; Massen et al., 2010). Finally, we expect an individual to provide a group member when they are peers have a positive relationship (a high relationship quality), or when the other has a higher rank in the dominance hierarchy (Massen et al., 2011; Massen et al., 2010).

## METHODS

### Subjects & Housing

The observations and behavioural experiments took place at the Biomedical Primate Research Centre (BPRC) in Rijswijk, The Netherlands. Three groups of long-tailed macaques (group I, N = 4; group II, N= 9 + 2 dependent female infants; and group III, N= 19) were tested (Table 1). Group I was a multimale group; Group II was a multigenerational group with an unrelated adult alpha-male, and group III was a multigenerational group without an unrelated adult alpha-male. Groups II and III lived in connected indoor (surface area: 73 m^2^, height: 2.85 m) and outdoor (surface area: 251 m^2^, height: 3.10 m) enclosures. Group 1 occupied a smaller indoor and outdoor enclosure, approximately 2/3 of the regular area.

**Table 1.** Characteristics of all individuals per group and their performance in all test trials, empty trials and blocked control trials. Dominance rank was calculated on the basis of submissive behaviours in dyadic encounters, age was calculated in months at the beginning of the first session. Pulls in test trials are the total number in 5 test sessions. Provisioning pulls are pulls in test trials when a food reward was provided and taken by a receiver.

| Group | Name | Abbr. | Dom. rank | Age (months) | Sex | Pulls in test trials | Pulls in control trials |  | Provisioning pulls |
| --- | --- | --- | --- | --- | --- | --- | --- | --- | --- |
|  |  |  |  |  |  |  | Empty | Blocked |  |
| I | Sollie | so | 1 | 61 | Male | 5 | 7 | 14 | 1 |
| I | Kluivers | kl | 2 | 39 | Male | 31 | 11 | 37 | 11 |
| I | Horsten | ho | 3 | 39 | Male | 24 | 4 | 11 | 8 |
| I | Driel | dr | 4 | 37 | Male | 0 | 0 | 0 | 0 |
| II | Lagoon | la | 1 | 77 | Male | 3 | 0 | 4 | 0 |
| II | Urbi | ur | 2 | 74 | Female | 8 | 0 | 11 | 3 |
| II | Tutudetuu | tu | 3 | 25 | Male | 28 | 10 | 20 | 12 |
| II | Tebbi | te | 4 | 40 | Female | 5 | 1 | 4 | 2 |
| II | Walibi | wa | 5 | 99 | Female | 10 | 1 | 8 | 4 |
| II | Toffi | to | 8 | 40 | Female | 32 | 0 | 3 | 16 |
| II | Mooi | mo | 9 | 40 | Female | 1 | 3 | 2 | 0 |
| II | Dukki | du | 10 | 38 | Female | 18 | 13 | 11 | 7 |
| II | Alibi | al | 11 | 39 | Female | 19 | 12 | 12 | 13 |
| III | Castello | ca | 1 | 158 | Female | 2 | 0 | 3 | 2 |
| III | Limoncello | lc | 2 | 25 | Male | 22 | 10 | 8 | 9 |
| III | Saboteur | sb | 3 | 163 | Female | 3 | 0 | 4 | 1 |
| III | Tamayo | ta | 4 | 41 | Female | 17 | 3 | 9 | 1 |
| III | Monopoly | mp | 5 | 21 | Male | 11 | 14 | 9 | 4 |
| III | Lageveen | lv | 6 | 29 | Female | 3 | 5 | 8 | 0 |
| III | Hacienda | ha | 7 | 158 | Female | 0 | 0 | 0 | 0 |
| III | Magelaen | mg | 8 | 163 | Female | 0 | 0 | 0 | 0 |
| III | Rox | ro | 9 | 46 | Female | 8 | 9 | 6 | 7 |
| III | Etten | et | 10 | 58 | Female | 0 | 5 | 2 | 0 |
| III | Freggel | fr | 11 | 10 | Male | 19 | 1 | 0 | 9 |
| III | Soest | ss | 12 | 70 | Female | 1 | 0 | 1 | 0 |
| III | Sokoban | sk | 13 | 158 | Female | 1 | 2 | 4 | 1 |
| III | Emerald | em | 14 | 23 | Female | 9 | 1 | 2 | 4 |
| III | Stas | st | 15 | 79 | Female | 0 | 3 | 1 | 0 |
| III | Manga | ma | 16 | 156 | Female | 0 | 1 | 1 | 0 |
| III | Aquileie | aq | 17 | 167 | Female | 0 | 0 | 0 | 0 |
| III | Nellie | ne | 18 | 21 | Female | 4 | 1 | 1 | 3 |
| III | Jakkiebak | jb | 19 | 169 | Female | 0 | 0 | 0 | 0 |

The animals were used to observers and had access to lots of enrichments and climbing opportunities (Vernes & Louwerse, 2010). They were fed twice a day, in the morning with monkey chow, in the afternoon with fruit or vegetables, and water was available ad libitum. Engaging in the behavioural experiments was on a voluntary basis, without food or water deprivation. Permission for the research was granted by the institute’s Animal Welfare Body (IvD: Number IvD-019) and complies with Dutch and EU law.

### Observational data collection

We observed the groups of long-tailed macaques to obtain information about the social relationships, using two different approaches: scan sampling (125 scans per group) and ad libitum observations (50 hours per group). We made scans at least 30 minutes apart with a maximum of 8 per day and noted the exact location of all visible individuals, describing the proximity to individuals within one arm’s length. These scans were conducted in the inside enclosure on the same days as we did the behavioural experiments.

The relationship quality of a dyad was measured based on the mutual proximity in the scans, a valid measure to distinguish between friends and non-friends (Massen, Sterck, & de Vos 2010). We summed up when two macaques were within one arm’s length of each other, divided by the total number of scans. When both individuals were not visible during a scan, we assumed that they were in proximity in the same ratio. The calculation is explained in more detail in table S4 of the Supplementary Data. In addition, for analysis of individual characteristics, we calculated general sociality per individual as the average relationship quality, by averaging their dyadic Z-score transformed proximity (see below) with all other group members.

The dominance hierarchy was determined by scoring all submissive behaviours seen during encounters. Unidirectional behavioural elements indicating submission, i.e. silent bare teeth, make room and give ground (cf. Angst, 1974), were scored ad libitum (Martin & Bateson, 2018). For each group a dominance hierarchy was calculated using Matman (Schmid & De Vries, 2013), a package that was performed in R (version 4.1.3). The ensuing dominance rank is provided in table 1, with number 1 being the highest-ranking individual per group. In both group II (linearity index h’ = 0.605; directional consistency index = 0.987; p = 0.0165) and group III (linearity index h’ = 0.457; directional consistency index = 0.970; p = 0.0005) the dominance hierarchy was significantly linear. For group I, we had only 4 animals, which did not allow us to test their dominance hierarchy statistically (Appleby, 1983), but their submissive behaviours were unidirectional and transitional. We based their dominance hierarchy on the unidirectionality of these observations.

Age and kinship were obtained from files of the BPRC colony management. To compare the age of individuals between different groups, we expressed age in months at the date the experiments started. Dyads with a relatedness of 0.5 (mother, brother, sister) or 0.25 (aunt, nephew, niece) according to the matrilines were considered kin.

### Experimental data collection

#### Test apparatus

The group service paradigm (cf. Burkart & Van Schaik 2013) was conducting by using an adjusted apparatus, the ‘swing set’ (Figure 1). This swing set contained a beam of 1.2m which hung from the ceiling by two steal chains. In the middle of the beam (position 0) and at the left-end side (position 1) a bucket was fastened to provide a small food reward. The swing set was attached in front of the enclosure, at a distance of 0.5m approximately, out of reach for the macaques inside. The only way they could reach for the swing set was to pull a rope, attached to the middle of the beam at position 0 and attached with a carbine hook to the fence. The swing set was positioned in such a way, that a macaque at position 0 could not reach for the bucket at position 1. Inside the enclosure a wall obstructed direct access; the monkey who pulled had to release the rope to walk through an opening in the wall to be in position 1. Gravity consequently drew the beam out of reach for the macaque that pulled the rope.

**Figure 1.**
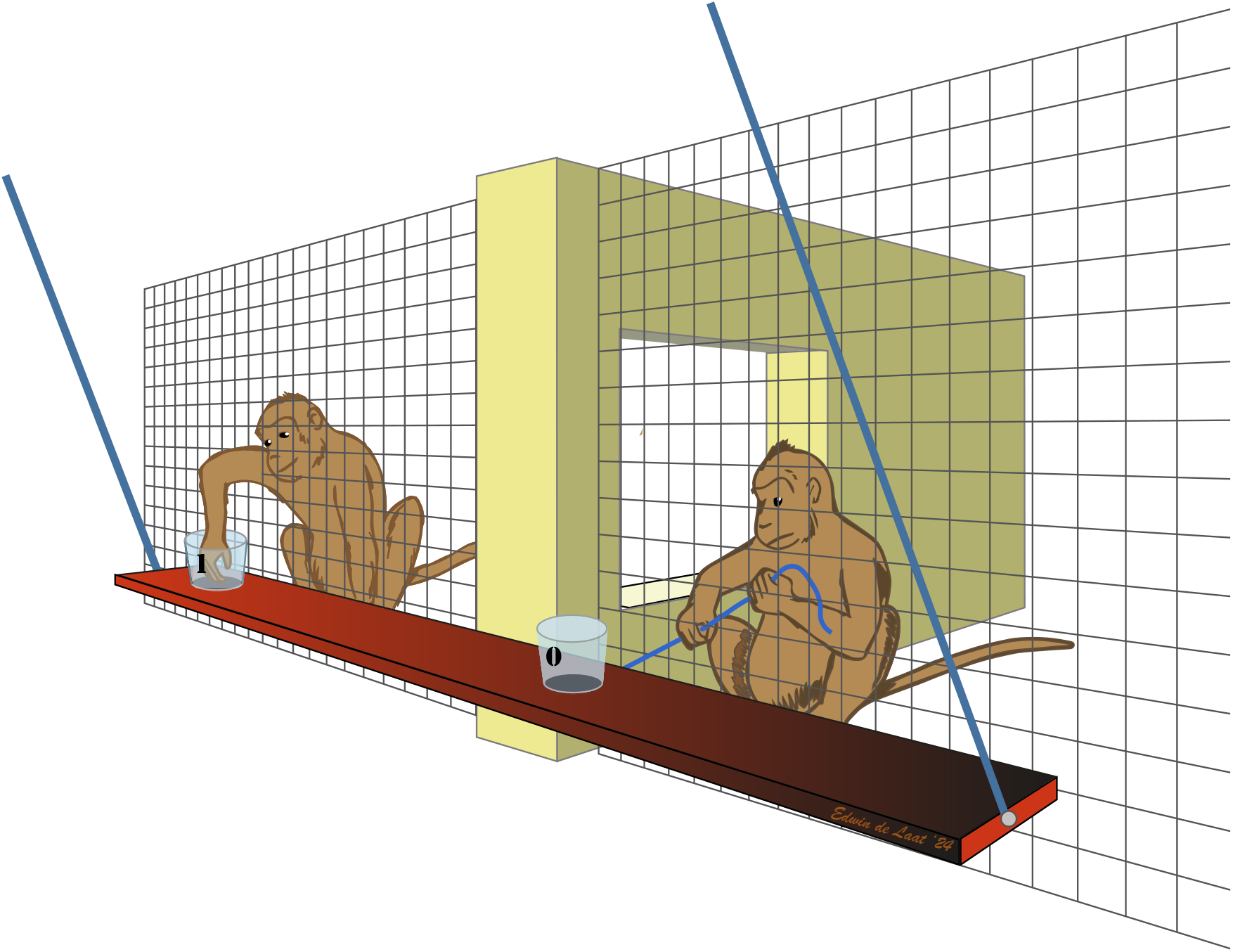
Swing Set Apparatus, the macaque in position 0 is providing a food reward to another individual in position 1. The macaques can see and interact with each other through an opening in the wall. The macaque in position 0 cannot reach for the bucket in position 1, because it has to let go of the rope.

### Experimental design

The experiment consists of five different phases (cf. Burkart & Van Schaik, 2013): a habituation phase (I); a phase to test for social tolerance (II); followed by a training phase (III); and a test phase (IV). After the regular tests and controls, we conducted a blocked control phase (V). Every session was video recorded with two cameras. We tested two times a week, in an indoor setting, under comparable circumstances in all three groups.

#### Phase I: Habituation

The beam was directly attached to the outside of the enclosure. All long-tailed macaques were able to reach into the bucket at position 0 to take a food reward (one piece of maize grain, pea or sunflower seed), during a session which lasted 90 minutes. When in one habitation session an animal had had 10 rewards, we ignored this experienced individual and tried to “invite” other individuals to the treats on the beam to get as many participants as possible. When every individual that took a reward passed the criterium of 10 successful attempts per session, we ended the habituation phase and continued to the next phase. In group I 3 out of 4 macaques succeeded, in group II 6 out of 9 and in group III 12 out of 19 macaques passed the criterium.

#### Phase II: Social Tolerance Test

In two separate sessions, on two different days, we tested the accessibility of the attached swing set for each individual in the different groups. While the beam was attached to the fence (cf. phase I), we baited the middle bucket (position 0) every 2 minutes during 5*n trials (n = number of individuals successfully participated in the habituating phase). We did two identical sessions in each group. The number of successful attempts to obtain food per individual was scored.

#### Phase III: Training

During the training phase, the beam could move like a swing, hanging from the ceiling outside of the enclosure, with a rope attached to the fence. If a monkey pulled the rope, the beam came forth and the monkey was able to take a reward out of the bucket in position 0. Our criterium to consider the training completed for a macaque in the group, was 10 pulls where it obtained a reward in a session. Again, if an individual had had pulled more than 10 times during one session, we ignored this individual and gave opportunities to others. We wanted to avoid monopolisation and aimed to achieve a maximum of individuals that participated. The next phase started when no new macaques participated voluntarily any further. To ensure that the animals understood the contingencies of the test apparatus, we performed sufficient training sessions in the different groups (Group I: 76 trials in 2 sessions; Group II: 177 trials in 4 sessions; and Group III: 130 trials in 3 sessions). The number of animals that pulled the rope at least 10 times in at least one session was in group I: 4/4 (100%), in group II: 5/9 (55%) and in group III: 13/19 (68%). In total 22 macaques passed our training criteria, which corresponds to 69% of all long-tailed macaques in this study.

#### Phase IV: Group Service Test

In this phase, during the regular test trials, we baited only the bucket at position 1 with a food reward (maize grain, pea or sunflower seed) and registered which individual pulled the rope in position 0 to move the swing set towards the fence. We also noted the individuals that were present or approached bucket 1 and whether they successfully obtained the food reward. To keep the participants motivated, we provided bucket 0 with a food reward in motivational trials. Every session started with a motivational trial, followed by five regular test trials. These six trials were repeated in each session as many times as we had participating macaques (n) in the habituation phase in that group. We ended every session with a motivational trial. In group I the number of participating macaques n=4 resulting in 25 trials; and in group II n=6 resulting in 37 trials. In group III n=12 participated, but due to BPRC welfare regulations that allowed a maximum duration of 90 minutes for a behavioural experiment, we had to limit the number of trials for this group to 43 trials of 2 minutes each. In every trial, we noted a pull when an individual pulled the rope so far, that the bucket in position 0 was accessible. We noted a provisioning pull when another individual took the reward in bucket 1. It was possible to have more than one pull in one regular trial; i.e. when no individual took the reward on the receiving end (see table S1 in the Supplementary data for the number of extra pulls in each session), but there could only be one provisioning pull, since there was only one reward in every regular test trial.

If three consecutive motivational trials were neglected, we would end the session. This occurred once in session 5 in Group III, because many individuals were attracted to something outdoors. We stopped this session after 31 trials and started all over again a day later. This incomplete session was ignored in our analyses and we treated the next, completed, session as session number 5. To be sure that the macaques understood the contingencies of the task; i.e. if they would keep pulling without rewarding themselves, we executed five sessions in total per test and control conditions (cf. Bhattacharjee et al., 2023; Burkart & Van Schaik, 2013). In the first three sessions, the results may be affected by learning attempts and some pulls may be unintentional. Therefore, in the analyses we only used the results from the last two sessions of these test trials and of the controls (see below)

To further ensure the monkeys understood the consequence of their behaviour, two types of controls were done. We alternated the regular test-sessions in Phase IV with empty control sessions, where we conducted the exact same experiment without a reward in position 1 in the empty control trials. By pretending to bait bucket 1, we drew attention to each trial and scored the responses of the macaques when there was no reward in bucket 1. In the empty control trials we also counted the pulls by the participating individuals, no provisioning pulls were possible, because rewards were absent. These empty control sessions consisted of the same number of trials, and, similar to the test trials, in the intermitted (once every 6 trials) motivational trials we baited bucket 0 to keep the macaques interested. In our calculations we compared the test results with the control results. The other type of control was conducted in Phase V.

The provisioning pulls in regular test trials resulted in providing another individual, the receiver, with a food reward. When a macaque, the provider, pulls in position 0 when a receiver is present in position 1, or close enough to reach for the reward, this was considered an act of reactive prosociality. When no individual was present in position 1 when the provider starts to pull, this pull was considered an act of proactive prosociality.

#### Phase V: Blocked Control

The second control experiment was a blocked control situation where a transparent screen, attached to the fence in front of position 1, prevented a receiver from taking the reward. The aim of this set up was to check if the provider would pull anyway, even if the screen made it impossible for the receivers to reach for the reward. We included both baited and empty blocked control sessions to make both phases comparable (cf. Bhattacharjee et al., 2023; Burkart & Van Schaik, 2013). We counted per session for each individual the number of pulls in the baited blocked control trials with a reward in bucket 1.

In phase IV and V, we saw some attempts by specific individuals to reach for the reward in bucket 1 by themselves (Phase IV: 6 ind. on average 15.33 attempts; range 8-26, with the majority of these pulls, i.e., 67,4%, in the first two sessions; Phase V: 2 ind. with both only 1 attempt). To obstruct these events, we pulled the swing set back (out of reach for the macaques inside the enclosure) with a rope attached to our side of the beam. However, when a macaque succeeded in its attempt to take the food reward in bucket 1, we omitted this trial from the dataset, i.e., we did not count this as a normal pull.

### Data analysis

We used R version 4.4.0 for statistical analysis of the data and to produce all graphs.

To determine tolerance in the social tolerance test - phase II - the evenness scores resulting in a Pielou’s J (Pielou, 1966) were determined for each test day and averaged. To measure the rate of participation of each individual, we counted the number of rewards they took. Pielou’s J ranges from 1.0, every macaque gets the same number of food rewards, to 0, one individual obtains all food rewards.

We compared the number of pulls during the regular test trials in the test sessions versus the empty control sessions of phase IV and the blocked control sessions of phase V to determine prosocial behaviour. We performed a Friedmans test to explore differences in number of pulls, and post-hoc a Wilcoxon test to distinguish between the conditions. For this comparison only sessions four and five were taken into account, to avoid an effect of learning.

Similarly, we compared the number of pulls in the final two sessions of the test and controls per individual, using chi—squared tests, to determine on an individual level whether an individual could be considered prosocial. Note here that the number of pulls each individual could perform in the (test) sessions was restricted by the number of pulls of all other participating individuals. To analyse the characteristics of prosocial individuals, we considered all individuals that pulled significantly more in the test sessions than in at least one of the controls as prosocial individuals (cf. Bhattacharjee et al., 2023; Horn et al., 2016; 2020). To analyse what characteristics, be it age, rank, sex, or general sociality had an effect on the likelihood of an individual to be prosocial, we ran a binomial GLM.

The combination of actor and receiver when food is provisioned is a key to unravel which characteristics determine a successful delivery of the reward. However, in many possible dyads we observed no prosociality, which made our data heavily zero-inflated. Our solution to this was fitting an implicit hurdle model to investigate (1) which dyadic factors predicted the occurrence of pulls within a dyad (binomial GLMM) and (2) which dyadic factors predicted the total number of pulls within dyads in which at least one pull had been observed (Poisson GLM).

To describe each dyad, we determined these characteristics: same sex (actor: male and receiver: male or both female) or different sex (actor: male and receiver: female or actor: female and receiver: male), age difference (age of actor in months minus age of receiver in months) and maternal kin or non-kin. Other factors we took into account were rank-difference and relationship quality. To standardise the rank-difference (rank actor minus rank receiver) and relationship quality in all groups, we z-transformed these data.

For the first model, we fitted a GLMM with a binomial error distribution using the lme4 package (Bates et al., 2015), with the occurrence of pulls within a dyad (yes/no) as dependent variable (N = 165). We included maternal kinship (yes/no, sum-to-zero coded), relationship quality (z-scored within group), age difference (z-scored overall), rank difference (z-scored within group) and the interaction between sex of actor (male/female, sum-to-zero coded) and receiver (male/female, sum-to-zero coded). Furthermore, we allowed the intercept to vary by Group and Actor and Receiver ID nested in Group. As sensitivity check, we also ran this model without data from group 1, because this group consisted only of non-kin males, thereby restricting their options to pull for kin or females. This model was entirely equivalent to the one described before, except that we could not include the random intercept at the group level.

In the second model, we used a GLM with a Poisson error distribution with the total number of pulls within each dyad as the response variable, regarding only the provisioning dyads (N = 29). We included the exact same predictors as in the first model. However, we did not have the opportunity to add random effects because of the low number of repeats per Actor and Receiver. In addition, due to diagnostic issues we could not fit the same model to the dataset excluding group 1 as a sensitivity analysis. Therefore, we opted to include only the significant main effects in the model excluding data from group 1.

For all models, we used the DHARMa-package (Hartig, 2022) to calculate scaled residuals to assess potential misspecifications. In addition, we calculated Variance Inflation Factos (VIF) to assess multicollinearity between predictors. We found no indications for model misspecification and all VIFs were below 5, indicating that no problematic collinearity was present. In the reported Poisson models, we also checked for overdispersion, but we found no indications of either over- or underdispersion. For the binomial models we report odds ratios and for the Poisson models we report incidence rate ratios, alongside z-values and p-values for each predictor. Furthermore, we first ran the models with interactions and tested these against a null model with only main effects using a likelihood ratio test. In case of non-significant interactions, we removed these from the model and interpreted the main effects from the simpler model (Engqvist, 2005).

## RESULTS

### Social Tolerance Test

Because of their despotic characterisation (Thierry et al., 2004), we predicted a great inequity in access to resources between the individuals of each group. However, in the Tolerance Test (phase II) in group I, 3 out of 4 macaques took at least one reward out of bucket 0. In group II 5 macaques out of a total of 9 participated successfully, and in group III we had 12 macaques from a group of 19 individuals, who took at least one reward from bucket 0. The calculated evenness score Pielou’s J was on average 0,63 (Group I: J’ = 0.65, Group II: J’ = 0.60 and Group III: J’ = 0.63), see for more details table S2 in the Supplementary Information.

### Group Service Test, group level

To test for prosocial preferences across our test animals, we compared the number of pulls in regular test trials during test sessions with the number of pulls during the empty control sessions and the blocked control sessions. When the provider would have the intention to reward another individual, it would not pull when there is no food reward present (in an empty control trial), or the food reward is not accessible (in a blocked control trial). During these test and control trials, three additional animals in group II and two in group III started participating, this renders our final sample size to 27.

The number of pulls in the regular test sessions increased and remained high, compared to the total number of pulls in both control sessions (Figure 2). The number of pulls in the control sessions seemed to decrease over time, suggesting that the individuals needed to learn the contingencies of these controls, whereas their prosocial preferences remained constant, even after repeated exposure to the fact that pulling in this test condition did not result in any reward for the provider itself.

**Figure 2.**
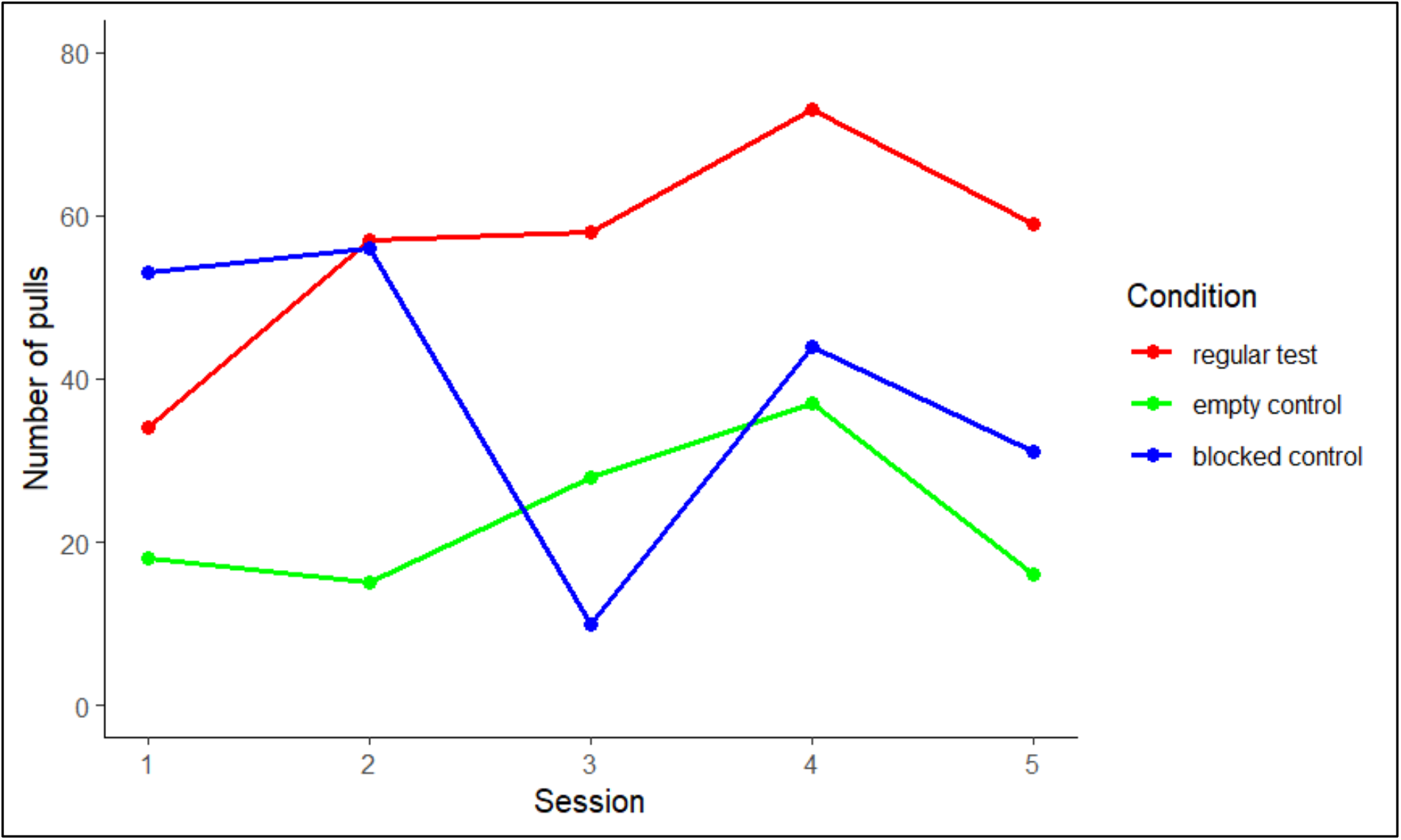
The total number of pulls in each of the five sessions in the regular test trials (phase IV), the empty control trials (phase IV) and the blocked control trials (phase V).

To ensure that in these trials all test animals understood the implications of both regular test and control sessions, the combined pulling of each individual in session 4 and 5 was compared. We found an overall significant difference in the number of pulls per condition (Friedman’s test: χ^2^ = 13.705, n = 26, df = 2, p = 0.001). Post-hoc comparisons revealed that the long-tailed macaques pulled significantly more often in the regular test sessions compared to both the empty-control sessions (Wilcoxon signed ranks test: p = 0.00046), and the blocked-control sessions (Wilcoxon signed ranks test: p = 0.04936). We found no difference between the two control conditions (Wilcoxon signed ranks test: p = 0.19765) (Figure 3).

**Figure 3.**
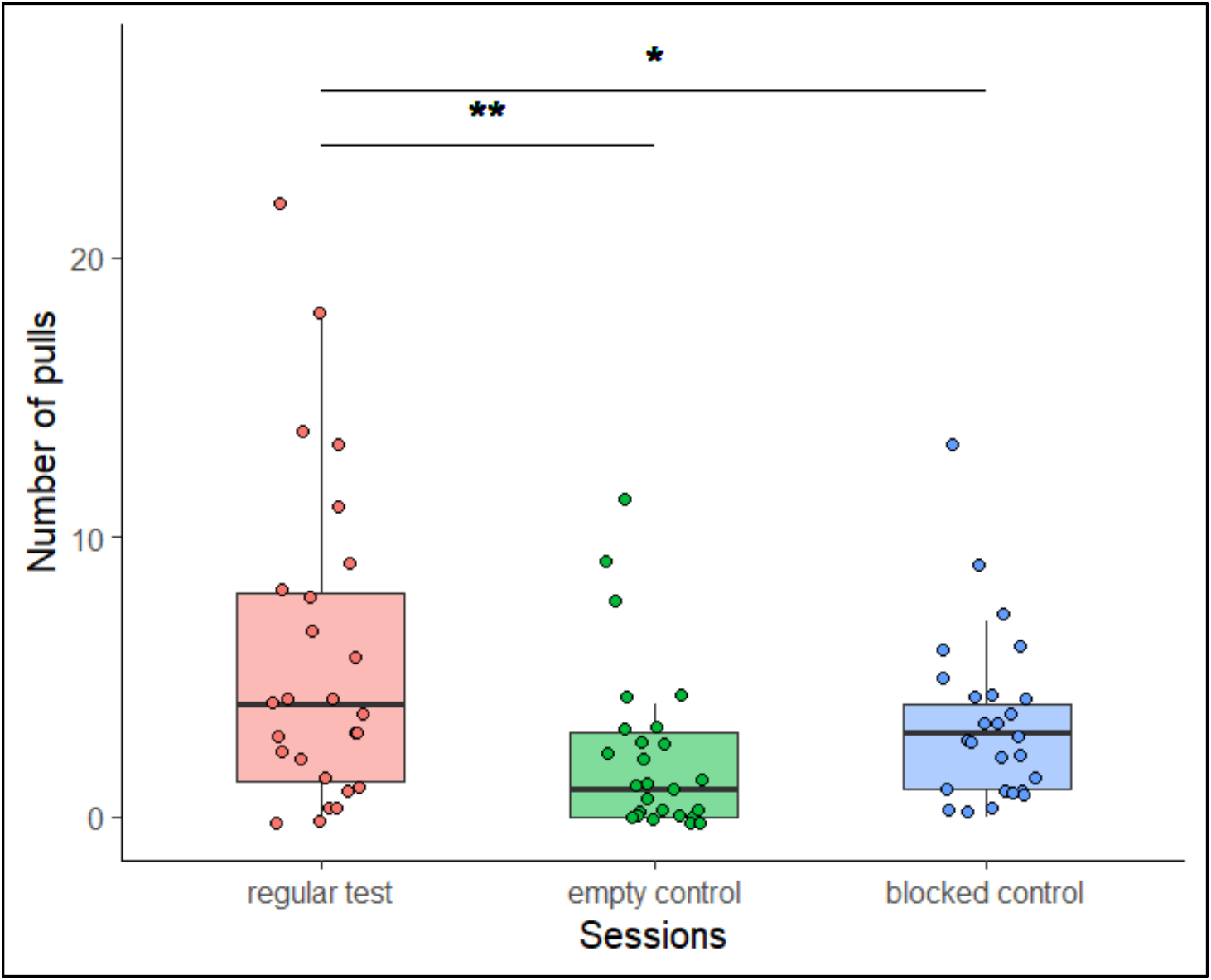
The number of pulls in the last two sessions (4 and 5) in the regular test condition (phase IV), the empty control condition (phase IV) and the blocked control condition (phase V). Dots represent individual data-points. Thick lines in box-plots represent the median, the box the 25 and 75-quartiles and the whiskers the 95% confidence intervals. * p < 0.05, ** p < 0.001

### Group Service Test, Individual level

We determined for each providing individual whether it was prosocial or not in sessions 4 and 5 (Table S3 - Supplementary Information). One individual, Horsten, pulled significantly more often in regular test sessions compared to both the empty and blocked control sessions (Test vs Empty Control: χ^2^ = 6.23, p = 0.013, Test vs Blocked Control: χ^2^ = 8.33, p = 0.004). Three additional individuals pulled more often in test sessions compared to the empty control (Kluivers: χ^2^ = 22.00, p = 0.000; Toffi: χ^2^ = 4.00, p = 0.046 and Emerald: χ^2^ = 5.00, p = 0.025) and three other individuals pulled more often in test sessions compared to the blocked control sessions (Alibi: χ^2^ = 9.80, p = 0.002, Dukki: χ^2^ = 5.00, p = 0.025 and Limoncello: χ^2^ = 5.00, p = 0.025). As the number of trials in the test conditions was limited and were further restricted by other pulling (and potentially prosocial) individuals, these comparisons can be rather conservative, and therefore, we considered these seven individuals prosocial (cf. Horn et al. 2016; Bhattacharjee et al. 2023). The other 19 individuals in the three groups did not show prosocial behaviour.

The contribution of the individual characteristics sex, age, dominance rank and average relationship quality, the general sociability on pulling in the regular test trials was determined (see table S4 in Supplementary Data for calculation, and table S5 in Supplementary Information for all characteristics). However, a GLM with the effect of the individual characteristics on the likelihood of these individuals to be prosocial or not revealed no significant effects (Supplementary Information: table S6).

### Group Service Test, Food provisioning

On average, the long-tailed macaques in the three tested groups pulled in 68% of the regular test trials (group I: 60 out of 100 trials; group II: 124 out of 150 trials; group III: 100 out of 165 trials). However, rewards were not always taken by group-members. If we consider trials in which there was a successful pull, on average only 28% of the regular test trials led to a successful food provisioning (group I: 20 out of 100 trials; group II: 57 out of 150 trials; group III: 41 out of 165 trials). Of these 118 providing pulls, 58 were proactive and 60 were reactive (reactive providing pulls: in group I: 11 out of 20 trials; group II: 21 out of 57 trials, group III: 28 out of 41 trials). If we then only consider those seven prosocial individuals that showed sufficient understanding of the task (cf. Burkart et al. 2013; 2014; Horn et al. 2016; 2020; Bhattacharjee et al., 2023), they are responsible for 58% (68 out of 118 trials) of the food providing pulls. When a receiver took a reward in position 1 that was provided by another macaque who pulled in position 0, it could potentially lead to some aggression by the provider towards the receiver. In all 118 pulls with a receiver, however, only 7 were followed by a threat from the provider. In 94% of the food provisioning interactions the provider allowed a receiver to take the reward. We never observed aggressive behaviour from the receiver to the provider during a trial.

### Dyadic predictors of provisional pulling

To investigate the dyadic predictors of provisional pulling, we performed an implicit hurdle analysis. As a first step, we investigated the likelihood of provisional pulling occurring for each potential dyad using a binomial GLMM (Table S7, Supplementary Data, for the dyadic characteristics and Table S8, Supplementary Information, for the results of the GLMM). We found no significant interaction between provider age and receiver age (χ^2^(1)= 0.70, p = 0.40). Therefore, we continued with a model containing only main effects. We found no significant effect of either age difference, rank difference, relationship quality, actor sex and receiver sex (Table S8a, Supplementary Information). However, we did find a significant effect of kinship (OR = 2.01, z = 2.35, p = 0.019), indicating that individuals were more likely to pull in kin dyads (Figure 4a). The model without data from group 1 (Table S8b) yielded the same result: only kinship had a significant effect on probability of observing provisional pulling within a dyad (OR = 2.01, z = 2.40, p = 0.017). To investigate which variables predicted the number of provisional pulls within dyads that had at least pulled once, we used a Poisson GLM. We again found no significant interaction between actor sex and receiver sex (χ^2^(1)= 0.42, p = 0.52). We found no significant effect of either age difference, rank difference, relationship quality, actor sex and receiver sex (Table S8c). However, we did find a significant effect of kinship again (IRR = 0.66, z = -3.208, p = 0.001), yet indicating that individuals pulled less frequently in kin dyads (Figure 4b). To make sure that these results were not caused by data from group 1, we ran a model on the dataset without group 1 with only kinship as predictor (Table S8d). This still resulted in a significant effect of kin in the same direction (IRR = 0.70, z = -2.507, p = 0.012).

**Figure 4.**
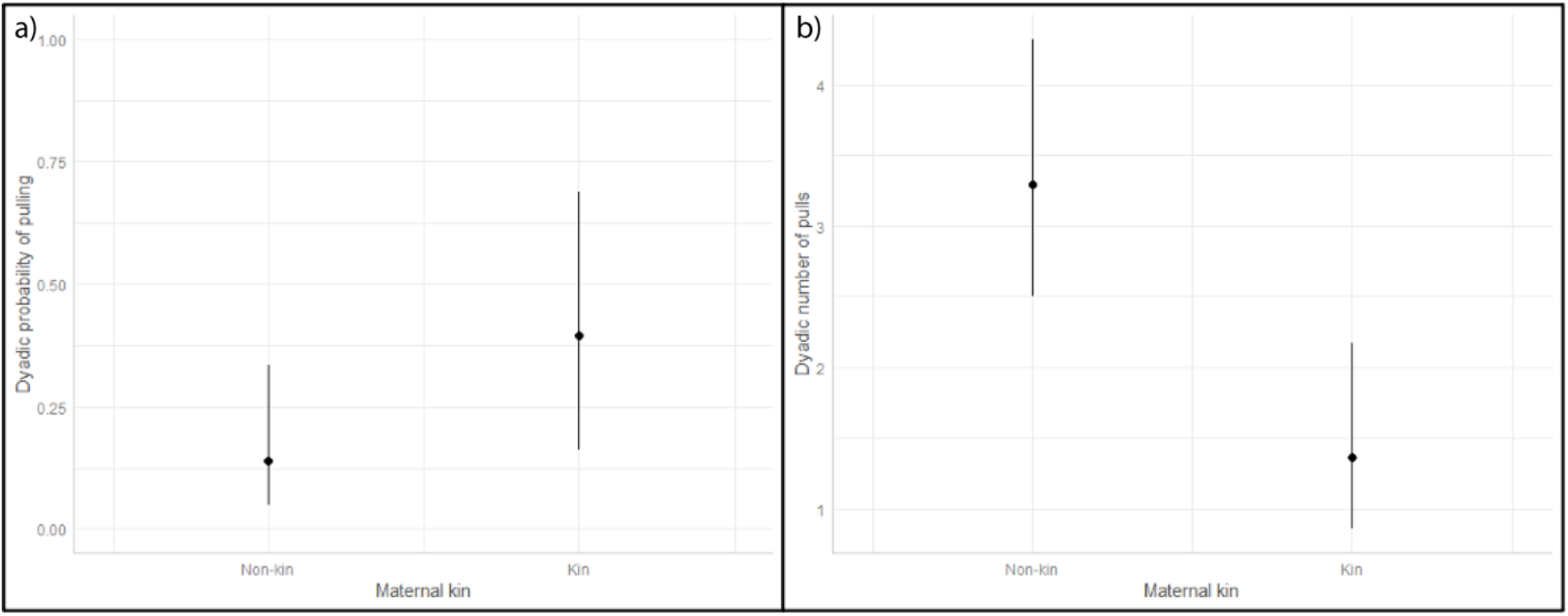
The a) dyadic probability of pulling and the b) dyadic number of pulls in non-kin and kin dyads. Dots represent the predicted value of the model, and whiskers the 95% confidence intervals

## DISCUSSION

Prosocial behaviour in a group service paradigm is expected in cooperatively breeding or self-domesticated primates (Burkart & Van Schaik, 2013; Jaeggi et al., 2010). Nevertheless, despotic macaques have also been shown to be prosocial in other prosociality tasks (long-tailed macaques: Massen et al. 2010; 2012) or in a group service paradigm (Japanese macaques: Bhattacharjee et al. 2023), possibly due to interdependency, i.e. when cooperators can benefit as a secondary consequence of helping their recipients (Roberts, 2005). Here we present evidence that long-tailed macaques, a species that does not breed cooperatively and is not egalitarian, showed relatively high tolerance at group level and acted prosocially in a group service task. Prosocial behaviour did not depend on individual characteristics of age, sex or dominance rank, yet provisional pulling was more likely for kin. Even in their unidirectional hierarchic organised groups, long-tailed macaques were rarely aggressive towards the beneficiary, when it took a food reward. Altogether, this supports earlier findings that prosociality is present in this despotic macaque species.

### Social tolerance

The first notable outcome of our study is the relatively high level of tolerance the long-tailed macaques exhibited. Many individuals participated in the tolerance tests and in all three tested groups we found a relatively high evenness score that is comparable to the score of the more egalitarian and food sharing brown capuchin monkeys (J’ = 0.66: (Burkart & Van Schaik, 2013)) and much higher than of the despotic Japanese macaques (J’ = 0.17: (Burkart & Van Schaik, 2013) and J’ = 0.13: (Bhattacharjee et al., 2023)), but lower than the evenness score of cooperatively breeding common marmosets (J’ = 0.74: (Burkart & Van Schaik, 2013). This relatively high tolerance may have allowed multiple individuals to handle the group service apparatus. Indeed, in the regular test and control trials in total 27 out of 32 individuals pulled the apparatus. The high tolerance is unexpected for a despotic macaque and contrasts with the low tolerance of Japanese macaques. In essence long-tailed macaques are considered a class less despotic than Japanese macaques (class II vs class I: Thierry. 2007), this may (partly) explain their higher level of tolerance.

Still, the long-tailed macaque dominance hierarchy typically explains the highly unidirectional type of aggression which can lead to differential resource access (Overduin - de Vries et al., 2020). In addition, long-tailed macaques have a unidirectional submission signal (i.e. the bared teeth display: Angst, 1974). Thus, the dominance relationships seem as obvious as in class I macaques. However, long-tailed macaques are considered class II due to their (slightly) higher rate of reconciliation, that may be possible due to relatively high tolerance for the opponent after a conflict (cf. Kempes et al., 2009). While aggression may result in access to resources, dominance position in itself does not (Overduin-de Vries et al. 2020), suggesting that less aggressive individuals may be more tolerant. Accordingly, in our study beneficiaries of the food rewards rarely received aggression from the provider (in only 7 out of 116 cases). Whether this difference in tolerance indeed distinguishes between despotic macaques of class I and class II, and of course is even higher in class III and VI (cf. Thierry, 2000; 2022), remains to be further explored.

### Prosocial behaviour

The long-tailed macaques were prosocial, both at individual and group level. They performed better in the test trials than in both control trials. This is congruent with earlier findings of prosociality in a dyadic or triadic setting in this species (Massen et al. 2010; 2011). In addition, this was found in each of the three groups, despite their different composition and size. This suggests that this is a general feature of this species and not a result of one outlier group. However, animals of this species were not prosocial in a dyadic setting when they had to forego a reward and choose the option without food to just provide someone else (Sterck et al. 2015). Yet, the results in the current group setting indicate that they can be prosocial in a less demanding setting where they receive no reward for providing another individual too.

The long-tailed macaques that participated in the test phase seemed to understand the task with the group service paradigm, also in our alternative design with a swing set apparatus, because in sessions four and five they pulled significantly more often when a food reward was available to a partner, compared to the phases when there was no food reward available (empty control) or when access was prevented (blocked control).

Our findings indicate that despotic long-tailed macaques, similar to Japanese macaques (Bhattacharjee et al., 2023), can be prosocial. This contradicts both the cooperative breeding and self-domesticated species hypotheses, but may support the alternative and overarching interdependency hypothesis (Bhattacharjee et al., 2023; Tomasello et al.,2012). This hypothesis argues that interdependency at either group or dyadic level or at both levels may lead to prosocial behaviour. While cooperative breeders may be dependent at the group level, in other group-living species interdependency may be found at the dyadic level, for example among kin or individuals with good relationships.

### Individual characteristics of prosocial individuals

Out of 27 participating individuals, 7 were significantly prosocial at the individual level. This is similar to the relative number of prosocial individuals in the despotic Japanese macaque (9 out of 25 individuals: Bhattacharjee et al., 2023). This is a conservative estimate of the proportion of prosocial individuals, since these prosocial individuals pulled relatively often and may have prevented other individuals from pulling and showing their prosocial behaviour at the individual level. In our experiment, the individual characteristics age, sex, dominance rank and general sociality did not affect whether an individual was prosocial. This contrasts with the earlier finding that dominants were more prosocial in a dyadic setting (Massen et al. 2010). However, in a triadic setting, subordinates were more likely to pull for dominant individuals (Massen et al 2011). In dyadic and triadic settings, prosocial behaviour always has a specific receiver, similar to the reactive setting in a group service paradigm. This suggests that the set-up of the experiment, i.e. dyadic, triadic or a group setting, may determine the importance of dominance rank in relation to prosocial behaviour, yet the current open setting, in combination with the restricted number of data-points we can gather from such an experiment, prevents analyses on the individual characteristics of prosocial individuals.

### Dyadic characteristics of prosocial behaviour

The interdependency hypothesis (Tomasello et al., 2012; Bhattacharjee et al., 2023) argues that interdependence at the dyadic level may promote prosocial behaviour in a despotic species like the long-tailed macaque. Indeed, when analysing behaviour during provisioning pulls, i.e. when someone obtained the food, we found that individuals were more likely to pull, at least once, for kin than for non-kin. This is consistent with the importance of kin for long-tailed macaques and fits the interdependency hypothesis. However, the number of provisioning pulls in prosocial dyads was less for kin than for non-kin individuals. This is an unexpected outcome that may be explained in several ways. First, subjects may need to be more prosocial to non-kin than to kin. To maintain a good relationship with kin may require less investment than a good relationship with non-kin. However, this was not borne out by our analyses, as we did not find an effect of relationship quality on the number of pulls. Second, some individuals may be more eager to obtain food, and be more often close to the provisioning location. This may, for example, concern individuals with a bold or persistent personality. Such persistent or bold individuals may obtain relatively large numbers of rewards, irrespective of who pulled. Unfortunately, we do not have data on personality. While this may explain the number of food items obtained, it does not explain that subjects were more likely to pull for kin. This line of reasoning suggests that the probability to pull for kin was not a by-product of personality. Third, receivers may be relatively dominant, keeping other individuals away from the provisioned location. Since individuals have more non-kin in the group than kin, when pulling is not directed at a specific individual, they will more often obtain food from non-kin than kin. A post-hoc analysis of the characteristics of the receivers (Table S7: Supplementary Data) showed more dominant individuals more often obtained a reward (Est. = -0.3994, z = -2.493; p = 0.0127), while sex, age and average relationship quality were not related to obtaining the reward. Thus, this reward taking by dominants may explain why we did not find that kin obtained more often rewards than non-kin.

Another interesting finding was that about half of the provisioning pulls were proactive, without a receiver present, and half were reactive, i.e. when a receiver was present in position 1. Similarly, in the despotic Japanese macaques 22% of the pulls were reactive, and majority (78%) were proactive, without a receiver near the baiting location (Bhattacharjee et al 2023). This indicates that also for long-tailed macaques, pulling was prosocial. Further exploring these two types of pulling may reveal whether reactive pulling is related to provisioning specific individuals, while proactive pulling is a prosocially motivated behaviour to the group in general (Jaeggi et al., 2010), that may be exploited by specific (e.g., bold, persistent, dominant) individuals. Unfortunately, for such analyses our data-set had to be split and these reduced data-sets rendered not enough statistical power to perform these analyses. Thus such analyses would require either a larger sample-size or more sessions of of the test condition.

Altogether, long-tailed macaques were more likely to pull for kin, although they pulled less often for kin than non-kin. This lower pulling for kin may result from dominants taking relatively often the provided reward. The preference to pull for kin is similar to long-tailed macaque behaviour in a dyadic paradigm (kin: Massen et al., 2010) and triadic paradigm (friends: Massen et al;, 2011). The pattern contrasts with the pattern in Japanese macaques in a group service paradigm, where dyadic tolerance predicted the likelihood of pulling and kinship predicted the number of pulls (Bhattacharjee et al., 2023). Nevertheless, this outcome is consistent with the prediction of the interdependency hypothesis (Tomasello et al. 2012; Bhattacharjee et al. 2023) that prosocial pulling is more likely found among dyadically interdependent individuals.

### Strengths and weaknesses

To compare prosocial behaviour in long-tailed macaques with other species, we conducted a group service paradigm (c.f. Bhattacharjee et al., 2023; Burkart & Van Schaik, 2013) in three different groups. The large number of participating individuals (N=27), the similar outcomes in the three groups and the many hours of observation make the results of this study robust. However, we could not use an apparatus with a pallung- (cf. Burkart and van Schaik, 2013), or see-saw mechanism (c.f. Bhattacharjee et al 2023) due to the facilities at the BPRC. A swing set was a solution with comparable characteristics: it swings back when a macaque lets go of the rope and handling the apparatus was easy to train.

Due to COVID-19 regulations at the BPRC, we were not allowed to come close to the enclosures (< 1m), therefore we baited the swing set apparatus with a stick. In the blocked phase trials we could not remove the rewards, which results in a piling of food items. This could have an exaggerated impulse for some macaques to keep trying to obtain the accumulating price. However, despite this caveat and the increasing number of rewards, the number of pulls was higher in the regular test trails than in the blocked control trials.

We did not perform a new pair of test sessions after the blocked phases (c.f. Bhattacharjee et al., 2023; Horn et al., 2016) to test the monkey’s understanding of the swing set apparatus. This was due to restrictions by the BPRC during the COVID-19 pandemic. Nevertheless, our results show an increase in the number of pulls during regular test sessions and a reduction in both types of control sessions, indicating that the long-tailed macaques understood the task.

### Conclusion

Prosocial behaviour in long-tailed macaques results in providing food rewards in a group service paradigm, although this is a despotic species with independent breeding that does not share food in a naturalistic environment. Consistent with the interdependency hypothesis, prosocial behaviour was more often directed towards kin, yet unexpectedly the monkeys pulled less often for kin than non-kin. However, this may be due to dominants obtaining relatively often the provided rewards. About half of the providing pulls was reactive, when a receiver was already present, the other half was proactive, when no-one was yet present. Thus, long-tailed macaques showed prosocial behaviour both for specific individuals (reactive) and to the group in general (proactive). Altogether, the prosocial behaviour of the despotic, individual breeding long-tailed macaques supports the dyadic interdependence hypothesis. Future studies on other macaque species with more tolerant dominance styles should elucidate the effect of dominance styles on prosociality.

## Acknowledgments

We thank Debottam Bhattacharjee for providing additional information on group I and Dian Zijlmans for her assist during data collection. Besides these collegues, we thank caretakers, veterinarians and staff of the BPRC for their dedication and professionalism which resulted in optimal housing conditions for the macaques.

